# LNISKS: Reference-free mutation identification for large and complex crop genomes

**DOI:** 10.1101/580829

**Authors:** Radosław Suchecki, Ajay Sandhu, Stéphane Deschamps, Victor Llaca, Petra Wolters, Nathan S. Watson-Haigh, Margaret Pallotta, Ryan Whitford, Ute Baumann

## Abstract

Mutation discovery is often key to the identification of genes responsible for major phenotypic traits. In the context of bulked segregant analysis, common reference-based computational approaches are not always suitable as they rely on a genome assembly which may be incomplete or highly divergent from the studied accession. Reference-free methods based on short sequences of length *k* (*k*-mers), such as NIKS, exploit redundancy of information across pools of recombinant genomes. Building on concepts from NIKS we introduce LNISKS, a mutation discovery method which is suited for large and repetitive crop genomes. In our experiments, it rapidly and with high confidence, identified mutations from over 700 Gbp of bread wheat genomic sequence data. LNISKS is publicly available at https://github.com/rsuchecki/LNISKS.

## Introduction

Bulk segregant analysis (BSA) involves pooling recombinant genomes to facilitate rapid identification of genetic markers associated with phenotypic traits (Michelmore *et al*., 1991; Giovannoni *et al*., 1991). Mapping-by-sequencing (MBS) combines BSA with second generation sequencing (SGS) to enable simultaneous mutation identification and mapping (Schneeberger *et al*., 2009). This original approach for identifying mutations in ethyl methanesulfonate (EMS) mutagenised populations relied on selection-induced patterns within genome-wide allele frequency (AF) in pooled genomes and was initially based on pooling 500 mutant F_2_ plants for sequencing. However back- or out-crossing of the mutagenised plants eliminates mutation load not linked to the causative mutation(s) as only offspring demonstrating the desired phenotype are retained (Zuryn *et al*., 2010). One such approach is MutMap where a mutant is crossed with the original wild-type followed by selfing of the offspring, which results in segregation of phenotypic differences in F_2_ progeny (Abe *et al*., 2012). Based on the assumption that the causative mutation occurs with highest frequency among bulked segregants a combination of isogenic BSA with deep candidate resequencing was applied to detect subtle allele frequency differences between closely linked mutations to facilitate the identification of causal ones (Hartwig *et al*., 2012). This technique can also be extended to identification of causal mutations from multiple independent mutagenesis events (Yan *et al*., 2017).

MBS is most powerful with whole genome sequencing (WGS) data but methods based on RNA and, more commonly, enrichment sequencing (e.g. exome capture) have been developed to address the issues of sequencing cost and computational challenges, particularly in the case of large and complex plant genomes (Gardiner *et al*., 2014; Mascher *et al*., 2014; Pankin *et al*., 2014; Ramirez-Gonzalez *et al*., 2015; Gardiner *et al*., 2016; van Esse *et al*., 2017; Wang *et al*., 2017). Alternative methods, such as MutChromSeq (Sánchez-Martín *et al*., 2016) and TACCA (Thind *et al*., 2017) rely on sequencing and assembly of flow sorted mutant chromosomes. Recently, AgRenSeq (Arora *et al*., 2018) was proposed as a powerful approach for detecting multiple disease resistance genes from crop wild relative diversity panels. AgRenSeq is particularly notable for its innovative way of linking phenotyping values to genotypic information represented by *k*-mers, which bares some resemblance to the HAWK approach used for disease association mapping in humans (Rahman *et al*., 2018).

A number of computational approaches have been developed for identifying mutations in MBS, mostly relying on a reference genome for aligning SGS reads (Candela *et al*., 2015). However, reference-based approaches are not always applicable or sufficient, typically due to lack of a suitable reference genome. The suitability of a reference genome depends on its completeness and level of conservation with the studied accession, which, particularly for large repetitive polyploids, needs to be high to allow reliable read alignment and subsequent variant calling. Considering the high diversity within globally cultivated crop species such as wheat (Jordan *et al*., 2015; Krasileva *et al*., 2017), there is no guarantee that the reference genome will be sufficiently similar to the studied variety in the chromosomal region of interest, e.g. due to presence of sequences introgressed from related species.

The established needle in the *k*-stack (NIKS) algorithm (Nordström *et al*., 2013) allows reference-free identification of homozygous mutations from WGS data. Briefly, two sets of *k*-mers are extracted from WGS reads. Set *W* contains *k*-mers from homozygous wild-type, and set *M* contains *k*-mers from homozygous mutant. A SNP can be represented by up to *k k*-mers in each of the two sets, these are the *k*-mers of interest. For example given the following sequence with a single base mutation: *ACG[***C**/**T***]TTA*, we identify three 3-mers {*CG***C**, *G***C***T*, **C***TT*} supporting the wild type allele and three 3-mers {*CG***T**, *G***T***T*, **T***TT*} supporting the mutant allele. The remaining 3-mers, namely {*ACG, TTA*} do not overlap the mutated base. Sets of sample-specific *k*-mers are identified through removal of *k*-mers which are present in both sets, that is: *W* ← *W* \ *M* and *M′* ← *M* \ *W*. Sample-specific *k*-mers from *W* and *M′* are unambiguously extended (assembled) separately, yielding sets *C*(*W′*) and *C*(*M′*) of contigs (or unitigs) which in NIKS nomenclature are called seeds. Of particular interest are contigs of length 2*k* – 1 which are likely to be centred around a mutated base, as there are up to *k k*-mers representing a SNP. Contigs from *C*(*W′*) are then paired with contigs from *C*(*M′*) to identify the mutations.

We have built on NIKS concepts to develop LNISKS (longer needle in a scanter *k*-stack, Figure 1), a high-throughput pipeline with a number of original features including a highly-parallelized assembly algorithm. We also introduce *k*-mer filters which can be generated from external data. The filters are expected not to contain *k*-mers matching those which support a putative causative mutation and so can be safely used to reduce the search-space and the incidence of false-positive calls. In addition, LNISKS addresses some of the challenges arising from uneven coverage common to SGS datasets through post-pairing extension of seeds under 2*k* – 1 bp. While NIKS has been shown to work in Arabidopsis (135 Mbp) and in Rice (430 Mbp) (Nordström *et al*., 2013), our approach scales to wheat-size genomes (17 Gbp).

**Figure 1:**
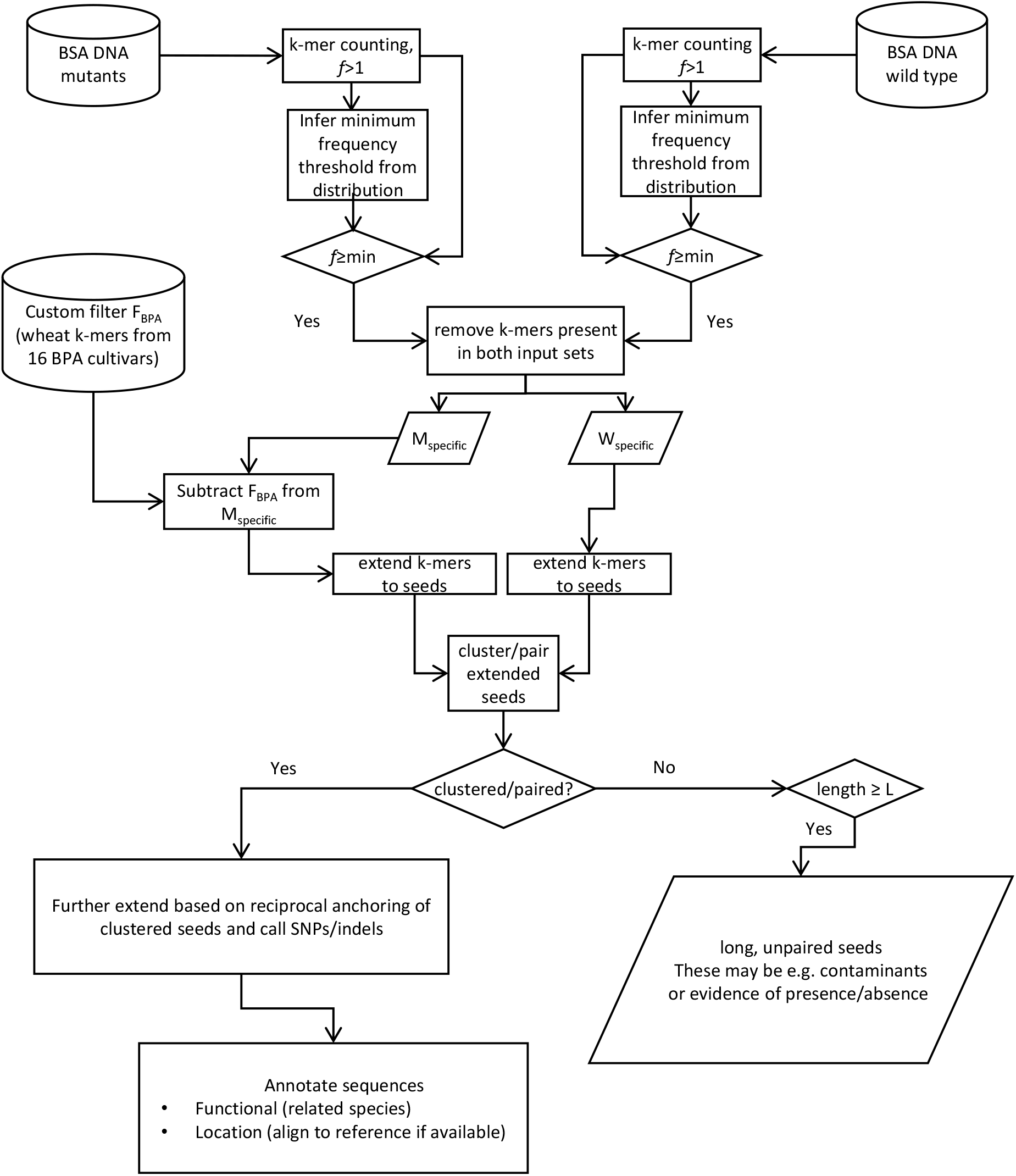
High-level overview of LNISKS approach.

Bread wheat genome is hexaploid and highly repetitive (Wicker *et al*., 2011; Choulet *et al*., 2014), so variant identification can be adversely affected by nearly-identical repeats and highly similar homeologous genes across the three closely related (sub-) genomes. In *k*-mer based approaches specificity can be improved e.g. by increasing k, potentially at the cost of reduced sensitivity. Longer *k*-mers are more likely to be unique within the genome, but require higher sequencing coverage to provide contiguous representation of the genome. As the number of *k*-mers in a genome increases with value of *k*, so does the computational cost of generating sets *W* and *M* from WGS, and comparing between them. We utilize KMC2/KMC3 (Deorowicz *et al*., 2015; Kokot *et al*., 2017), which allows fast, memory-efficient *k*-mer counting for *k* up to 256 and equally importantly, database level operations on sets of *k*-mers – most pertinently subtraction. A customized version of KMC3 has recently been shown to speed-up the early stages of the NIKS pipeline while reducing its memory requirements (*Kokot et al*., 2017).

We have used LNISKS to identify a mutation underlying *ms5* genic male sterility in bread wheat (Pallotta *et al*., 2019). In addition to a causative SNP, LNISKS also identified mutations underlying the markers which contributed to narrowing the *Ms5/ms5* critical region. The WGS data underlying these results comes from 20 *Ms5* (wild-type) and 40 *ms5* (mutant) plants. The mutant and-wild type bulks were generated from a cross of ms5 mutant plants with a male-fertile sib and the resulting F_2_’s were screened for homozygosity at the *TaMs5-A* locus using 5 markers. The combined genome coverage was ≈ 19X and ≈ 23X for wild-type and mutant bulks respectively. Based on these datasets we demonstrate the utility of LNISKS and explore some of its parameter space to shed light on capabilities and limitations of our approach.

## Results

We assess the accuracy of our pipeline and explore the effects of the innovations introduced in LNISKS. The first of our performance measures relies on the fact that we are looking for ethyl methanesulfonate (EMS) mutations, which are expected to be overwhelmingly G/C to A/T transitions (*Greene et al*., 2003). While exploring the parameter space the proportion of such transitions among our calls serves as a key benchmark of our pipeline’s accuracy. Another measure we employ relates to the expected lengths of seeds (contigs) extended from sets of sample-specific *k*-mers and consequently, to the paired/clustered seeds alignment length. A pair of contigs of length 2*k* – 1 bp with a single nucleotide polymorphism in the centre is the expected ideal for yielding a high-confidence variant call, as each of the two contigs is composed of all *k k*-mers which overlap either the wild-type or the mutated base, respectively. We classify such calls in the highest confidence category A. For a number of reasons however, contigs may either be shorter or longer than 2*k* – 1 bp. Pairs where variant position is *k* bases from one of the ends may also be of interest. We place calls corresponding to these pairs in confidence category B. All the remaining calls are assigned to category C as most likely to be spurious. The additional category D covers a subset of category B calls with more than one varying site per pair of sequences and at least one of these being *k* bases from a contig/alignment end. It covers the rare cases of two or more mutations within *k* bases from one another.

### The choice of *k*

Longer *k*-mers are more likely to be unique within a genome. Analysis of *k*-mer frequency plots (Figure 2) for *Ms5/ms5* data suggests that the longest *k*-mer for which the distribution does not appear to be truncated for either of the two input datasets is *k* ≈ 56. Higher *k* values in combination with limited sequencing coverage available result in an increased proportion of the target sequence not being captured by *k*-mers. Therefore, we would set *k* at or slightly below that value for further analysis. For illustrative purposes we explore a range of *k* values, *k* ∈ {24, 32,…, 72} and note that unless explicitly stated, the presented results pertain to LNISKS run at *k* = 54. The choice of *k* affects mainly accuracy but to some extent also computational requirements.

**Figure 2:**
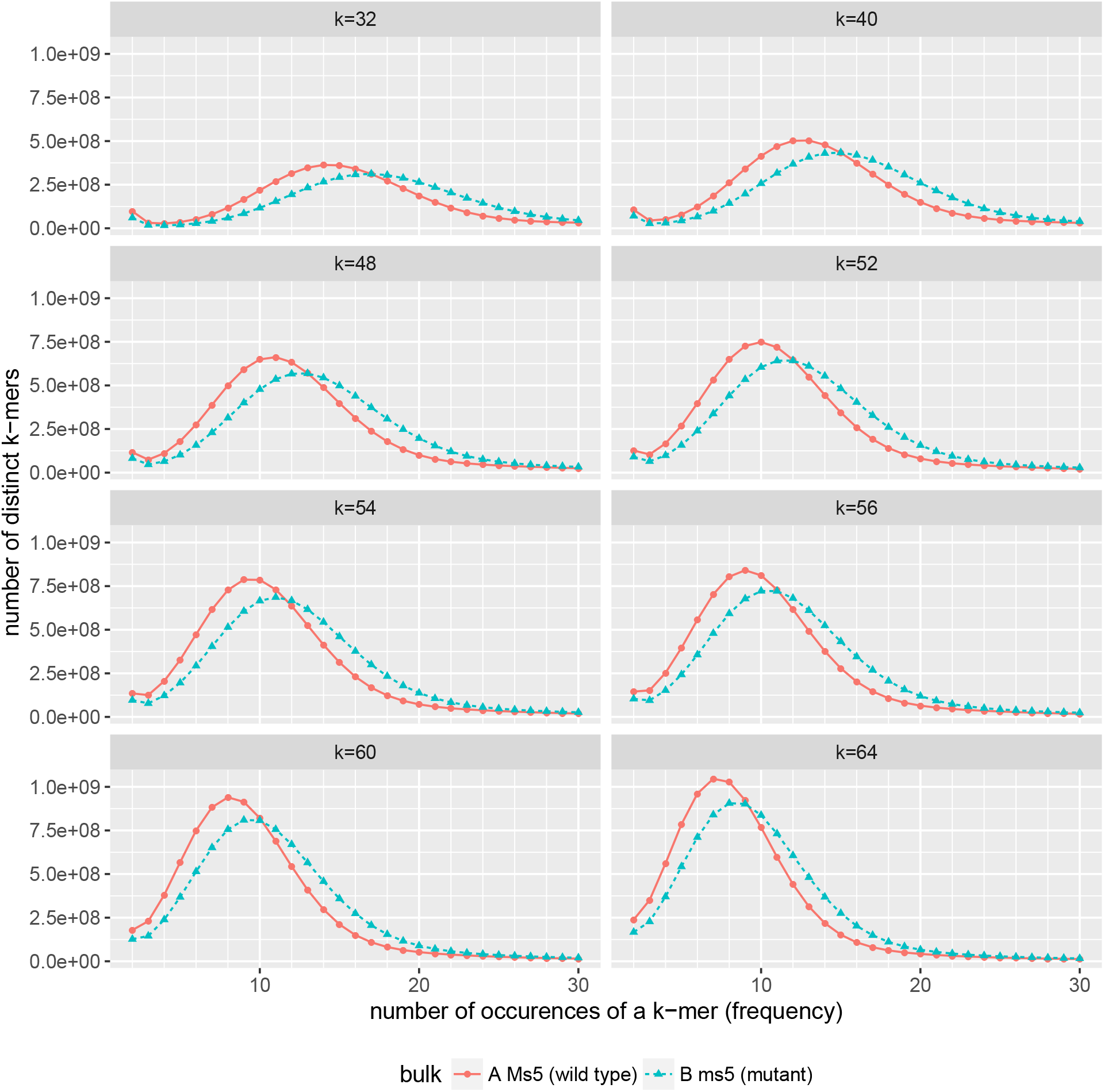
With increasing *k*, the number of distinct *k*-mers increases, but the frequency distribution shifts to the left, eventually becomes truncated and no longer captures the inflection point at frequency ≈ 3. This indicates insufficient sequencing coverage for contiguous assembly of mutation harbouring sequences at or above *k* ≈ 60. We can also observe that at low values of *k* the number of distinct *k*-mers is relatively low. This indicates that such short *k*-mers are unlikely to deliver sufficient specificity. Note that counts of *k*-mers of frequency 1 are not reported as these primarily reflect sequencing errors.

### Application of *k*-mer filters removes non-EMS-derived calls

Recall that the crucial step in the LNISKS (and NIKS) approach is the identification of the two sets of *k*-mers which appear in only one of the two bulks (typically wild-type and mutant). In LNISKS, once we obtain sets *W′* and *M′* of such sample-specific *k*-mers, we apply a novel *k*-mer filtering step, which is subject to availability of suitable data and specific biological context of the input datasets. Our experiments show that this step greatly reduces the number of *k*-mers considered for the assembly. This eases the computational requirements and helps to reduce the number of candidate mutations by discarding loci which may be regarded as irrelevant due to their presence in genomes which do not produce a given phenotype. As illustrated by Table 1, filtering reduces the number of calls to be considered/validated. The percentage of G/C to A/T transitions also indicates that the filtered-out calls are overwhelmingly not EMS-derived. Across the explored values of k, G/C to A/T transitions constitute about two thirds of category A SNPs, compared to around one third for other categories. If we apply our custom filtering step, the proportion of G/C to A/T transitions among category A calls increases to over 95% (Table 1), thereby approaching the level expected for EMS mutations (Greene *et al*., 2003). The number of category A calls and the number of G/C to A/T transitions called from category A clusters is highest at *k* ≈ 54 (Table 1) which is close to the choice of *k* that could be made based on the preliminary analysis of *k*-mer distributions for a range of *k* values (Figure 2), as described above.

**Table 1:** Numbers of paired seed for a range of *k* values. Each pair of seeds supports one or more putative mutations. Assignment to a confidence category A, B or C is based on the length of the alignment and the position of called mutation(s) within it (see text for details). The subscript _filtered_ indicates results obtained from runs where custom filters were applied to the set of *k*-mers from the mutant pool as described in experimental procedures. The bottom row indicates the percentage G/C to A/T transitions which are expected to dominate among true calls. These are largely consistent across the range of *k* values so we do not provide a per *k* breakdown.

| k | Number of paired seeds per confidence category |  |  |  |  |  |  |  |  |  |
| --- | --- | --- | --- | --- | --- | --- | --- | --- | --- | --- |
|  | All | All<sub>filtered</sub> | A | A<sub>filtered</sub> | B | B<sub>filtered</sub> | C | C<sub>filtered</sub> | D | D<sub>filtered</sub> |
| 24 | 12,652 | 4,792 | 2,758 | 1,579 | 7,160 | 2,710 | 2,734 | 505 | 249 | 16 |
| 32 | 17,192 | 7,079 | 4,629 | 2,753 | 9,297 | 3,630 | 3,266 | 698 | 431 | 32 |
| 40 | 28,113 | 13,176 | 5,848 | 3,824 | 15,419 | 6,702 | 6,846 | 2,652 | 529 | 50 |
| 48 | 34,416 | 15,170 | 7,155 | 4,679 | 18,855 | 7,392 | 8,406 | 3,101 | 813 | 76 |
| 52 | 34,561 | 14,955 | 7,678 | 5,005 | 19,785 | 7,405 | 7,098 | 2,547 | 483 | 56 |
| 54 | 36,747 | 15,789 | 7,766 | **5,228** | 21,547 | 7,861 | 7,434 | 2,702 | 609 | 58 |
| 56 | 51,539 | 21,770 | 7,656 | 4,967 | 30,805 | 11,831 | 13,078 | 4,974 | 689 | 58 |
| 58 | 80,229 | 42,908 | 7,343 | 4,276 | 44,788 | 22,926 | 28,098 | 15,708 | 671 | 64 |
| 60 | 83,342 | 44,681 | 7,379 | 4,455 | 47,372 | 23,901 | 28,591 | 16,327 | 725 | 82 |
| 64 | 95,873 | 50,790 | 6,540 | 4,029 | 53,250 | 26,769 | 36,083 | 19,994 | 811 | 126 |
| 72 | 63,532 | 59,557 | 4,381 | 4,641 | 38,083 | 32,817 | 21,068 | 22,101 | 213 | 122 |
| G/C to A/T | 36.1% | 64.3% | 67.5% | **95.3%** | 34.0% | 61.9% | 26.8% | 52.9% | 37.2% | 68.0% |

### Identification of mutations linked to TaMs5-A

Identification of the gene underlying the *TaMs5* locus relied on several bi-parental mapping populations, varied genomic and transcriptomic datasets as well as a range of bioinformatic techniques employed to generate relevant molecular markers for mapping the causative locus (*Pallotta et al*., 2019). Mutations between *Ms5* wild-type and the *ms5* mutant detected by LNISKS underlie many of the molecular markers contributing to that effort. The overall numbers of mutations detected are summarized in Table 1. The detected SNPs include a non-synonymous mutation in a gene demonstrated to be causative for *ms5* sterility (Pal*lotta et al*., 2019). It is found among putative mutations reported by our pipeline for the explored values of *k* ∈ {24, 32,40,48, 52, 56, 64, 72} irrespective of whether filters have been applied. This should not however be treated as an indication that the choice of the value of *k* or the filters do not matter, although the number of marker SNPs detected was similar at various settings. This may reflect the fact that sequences which are more unique across the genome were more suitable for use as markers and are also more easily detectable from BSA data. Note that the application of custom filters is of little relevance when identifying SNPs for molecular markers as the presence or otherwise of a SNP in other cultivars need not affect its suitability for that purpose. This is echoed in the negligible influence of the use of filters on the number of detected SNPs which were ultimately used as molecular markers (see Table 2).

**Table 2:** Number of key confidence category A markers identified at varying *k* values, with or without the application of custom *k*-mer filters.

| k | 24 | 32 | 40 | 48 | 52 | 54 | 56 | 64 | 72 |
| --- | --- | --- | --- | --- | --- | --- | --- | --- | --- |
| unfiltered | 17 | 18 | 22 | 21 | 20 | 19 | 20 | 18 | 18 |
| filtered | 17 | 18 | 22 | 22 | 21 | 20 | 20 | 19 | 17 |

### Evaluation of mutation calls using a reference genome assembly

With more complete and contiguous wheat assemblies becoming available (Clavijo *et al*., 2017; Zimin *et al*., 2017; IWGSC, 2018), we are able to use these to evaluate a significant proportion, though certainly not all, called variants. This limited applicability is to be expected, as the assemblies are for Chinese Spring, a variety distinct from Chris which is *ms5* background. The reference based evaluation is two-fold. We first focus on false positive calls which are a major issue for many reference-free approaches (Leggett and MacLean, 2014). This can be done using any one of the three aforementioned assemblies. We align paired seeds to the IWGSC RefSeq v1.0 (IWGSC, 2018) genome assembly to identify false positive calls postulated by those pairings. If each of the sequences from a given pair aligns to a different chromosomal location, the pairing is almost certainly spurious and so is the associated call. The results of this evaluation are summarized in Table 3. We were able to unambiguously align both seeds from over 90% of pairs which support category A calls. Among these, less than 0.5% were shown to be false-positive based on conflicting alignment locations. If custom *k*-mer filtering is applied, almost 95% of pairs unambiguously align and the associated false-positive rate falls to 0.1%.

**Table 3:** Paired contigs for *k* = 53 were aligned to the reference genome allowing 1bp indels and up to 3 mismatches per 100bp. Orphaned alignment of just one of the elements of a pair indicates that the other element did not align or that it aligned equally well at multiple locations. Although the SNPs represented by the multi-aligned sequences cannot be easily classified as false positive (FP) they are dubious at best. If both sequences from a pair align unambiguously to the reference, we compare their alignment positions and easily identify FP calls wherever the two contigs align to distinct locations in the reference genome.

|  | A | A<sub>filtered</sub> | B | B<sub>filtered</sub> | C | C<sub>filtered</sub> |
| --- | --- | --- | --- | --- | --- | --- |
| Input (pairs) | 7,742 | 4,418 | 20,661 | 6,837 | 7,352 | 2,544 |
| Orphaned | 410 | 89 | 5,152 | 1,541 | 2,538 | 901 |
| Both aligned (placed) | 7,032 | 4,177 | 10,146 | 3,217 | 1,427 | 339 |
| Both aligned (percentage) | 90.8% | 94.5% | 49.1% | 47.1% | 19.4% | 13.3% |
| Matched position | 7,000 | 4,173 | 9,465 | 3,091 | 873 | 214 |
| Identified False Positives (IFP) | 32 | 4 | 681 | 126 | 554 | 125 |
| IFP as percentage of placed | 0.46% | 0.10% | 6.7% | 3.9% | 38.8% | 36.9% |
| IFP as percentage of input | 0.41% | 0.09% | 3.3% | 1.8% | 7.5% | 4.9% |

The second reference-based evaluation relies on *a priori* information about the expected physical location of the locus of interest. Back-crossing is expected to remove EMS-derived mutations from the genomes of the individuals, except for regions linked to the locus causing the phenotype. We expect a concentration of SNPs in the linked regions. For *ms5*, this should be in the centromeric region of chromosome 3A. This evaluation is made possible by the IWGSC RefSeq v1.0 assembly (IWGSC, 2018) which allows investigation of the detected mutation load across the pseudo-chromosomes which incorporate an overwhelming majority of the assembled sequences. As illustrated by Figure 3, the G/C to A/T transitions are concentrated on chromosome 3A with certain blocks, primarily also on chromosome 3A, marked with presence of other mutations which are unlikely to be EMS-derived. These could indicate allelic variation associated with the *ms5* background Chris, and the majority of these are discarded if we apply our custom filtering step. We also observe a band of G/C to A/T mutations at ≈ 550 Mbp. This location corresponds to a super-scaffold in IWGSC RefSeq v1.0 which is most likely incorrectly placed within the pseudo-chromosome. Syntenic ordering of proteins along pseudo-chromosme 3A against pseudo-chromosome 3B as well as related species suggest that the super-scaffold should be placed at ≈ 100 Mbp point.

**Figure 3:**
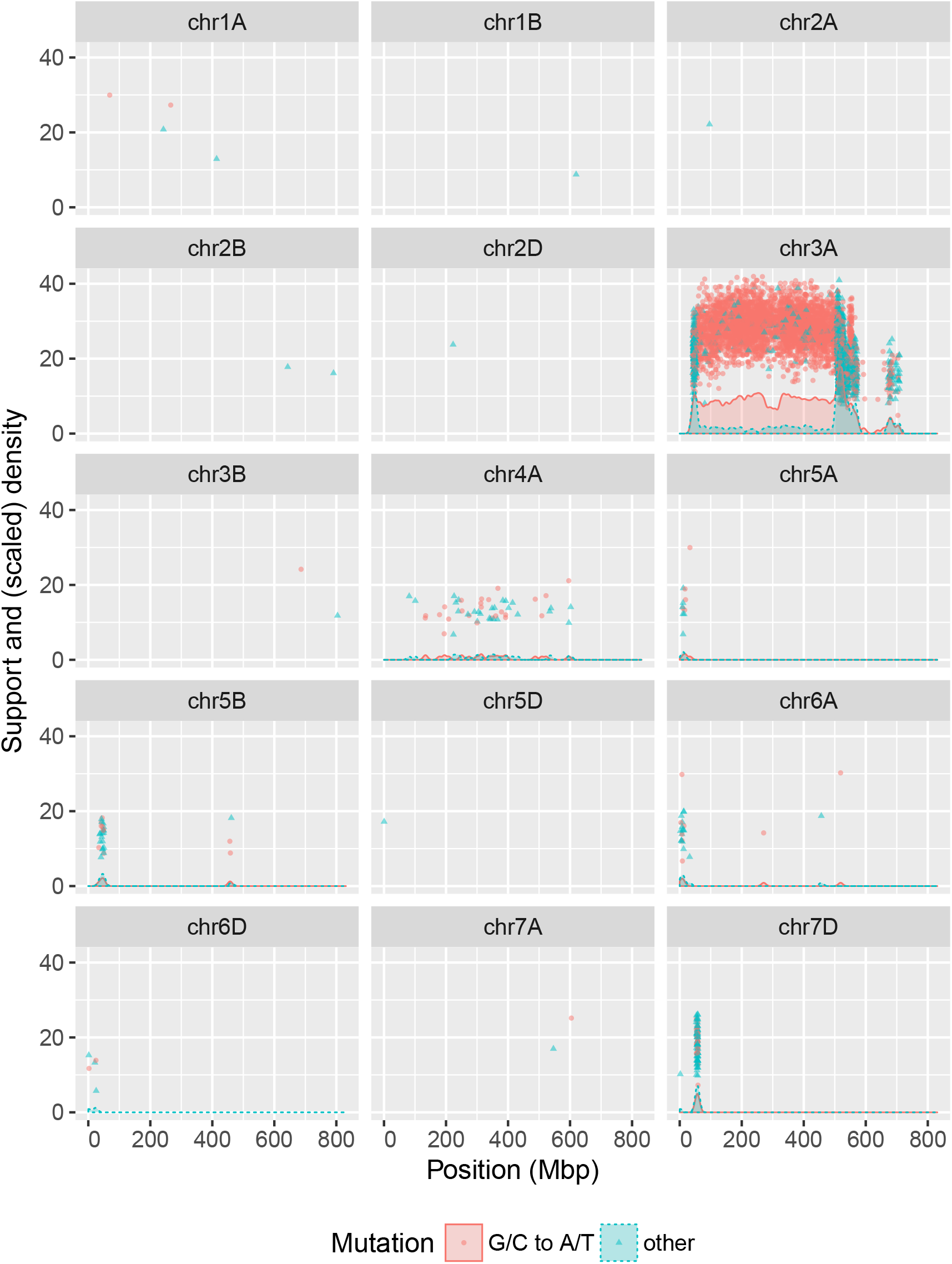
Confidence category A mutations identified without the application of filters, distributed along pseudo-chromosomes. Overall, 81.7% (83.7% filtered) of category A calls and 89% (92.8% filtered) of G/C to A/T transitions assigned to a chromosomal position are concentrated on chromosome 3A. Pseudochromosomes with no putative mutations assigned are not shown. Support for a given call is a simple measure calculated based on presence of *k*-mers supporting wild-type (mutated) allele in wild-type (mutant) plants. It is a crude reflection of the number of plants contributing evidence for a given call. The coloured areas under curves represent density of mutations within 5Mbp bandwidth. To aid visualisation (Wickham, 2009), the density estimate values are scaled as follows 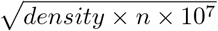, where *n* is the number of points.

When we look at chromosome 3A in more detail (Figure 4), we observe a large block (40 Mbp - 500Mbp) where many of G/C to A/T transitions display high support values, which reflect the number of plants contributing the underlying *k*-mer information. Furthermore, we observe how the application of custom filters discards many non-EMS derived SNPs, often concentrated just outside the large block rich in G/C to A/T transitions.

**Figure 4:**
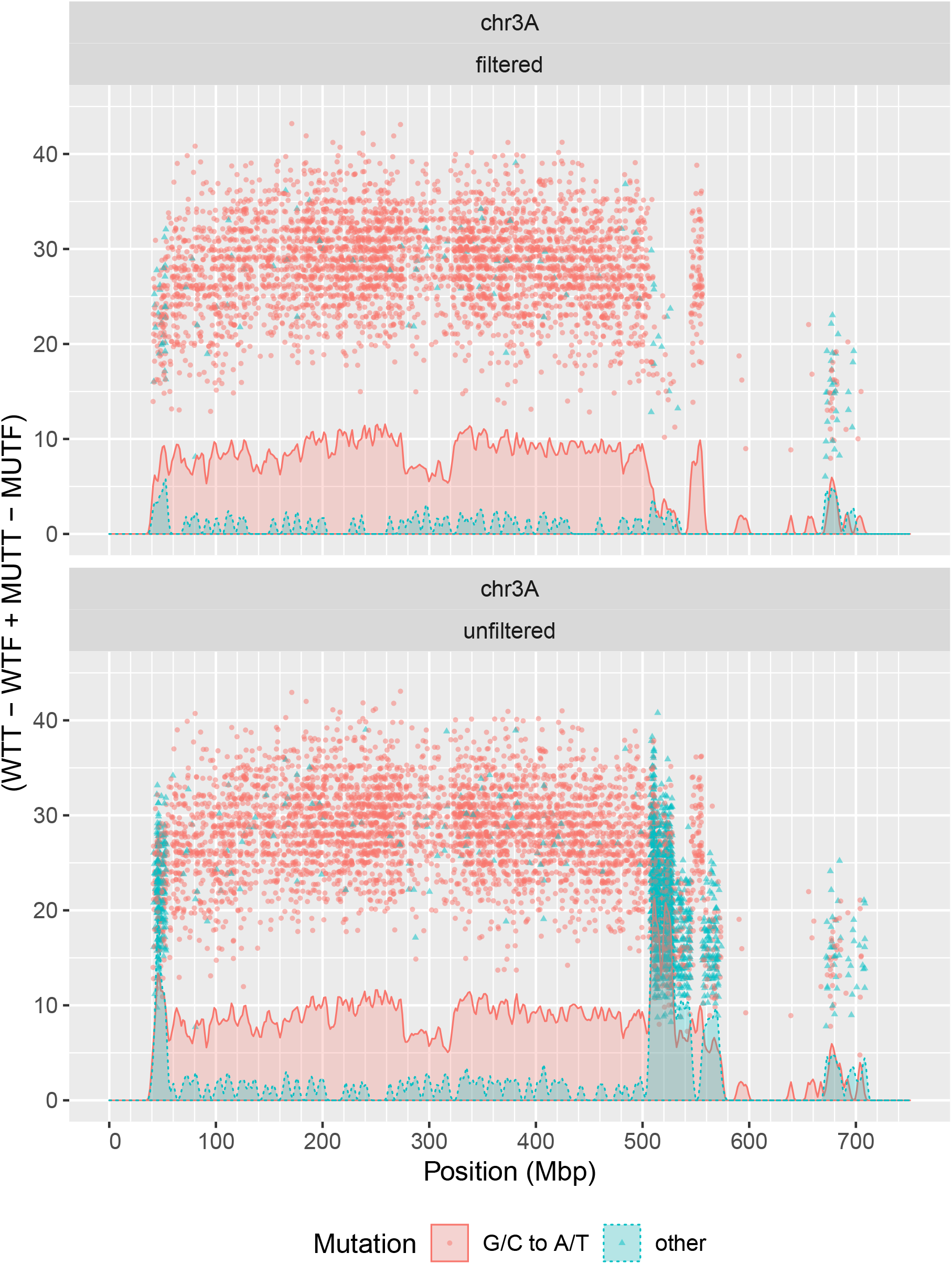
Confidence category A mutations distributed along the 3A pseudo-chromosome. Position of an individual data point along the y axis reflects the number of plants contributing *k*-mers which support the corresponding mutation. The coloured areas represent density of mutations within 1Mbp bandwidth. To aid visualisation, the density estimate values are scaled as follows 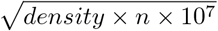, where *n* is the number of points.

### Computational cost

Through the use of state-of-the-art tools such as KMC2/3 and VSEARCH (Rognes *et al*., 2016) as well as extensive use of multi threading, LNISKS can process datasets consisting of billions of reads within hours. All *in silico* experiments were executed on an allocation of 32G of RAM and 16 logical cores on a compute cluster containing two nodes, each with 72 Intel Xeon E5-2699 v3 CPUs (2.30GHz), 770 Gigabytes RAM and two RAID0 SSDs for temporary files. Typical wall-clock run time of the pipeline (*k* = 53) was 2 hours and 33 minutes, which compares favourably with the 2 hours required for sequential, single-threaded reading and decompression of the same input data. The total CPU time for this run was 26 hours and 34 minutes. Where applicable, additional time is required for pre-computing custom filter database(s). For lower *k* values it may also be necessary to further extend paired seeds to facilitate better (sequence similarity based) functional annotation of contigs underlying putative mutations.

### Mutation identification from heterozygous data - proof-of-concept

Both NIKS and LNISKS are designed for detecting homozygous mutations. This enables straightforward subtraction of *k*-mers which in turn makes the approaches computationally tractable. This is a direct consequence of the fact that an overwhelming majority of *k*-mers are discarded by the subtraction step and the subsequent operations are carried in a much reduced search space. Here we present a proof-of-concept approach, where with the tool set comprised of KMC2/3, our custom filters as well as our vclusters and seedmers modules, we are able to quickly identify the *ms5* causative mutation and some of the key marker-SNPs for the locus when one of the input datasets (*Ms5* fertile) comes from plants which are phenotypically indistinguishable from the homozygous wild-type but known to be heterozygous based on marker information. This (third) bulk is comprised of 20 individuals, and the estimated combined sequencing depth was just under 15X assuming 17 Gbp genome. In this analysis we also use the bulk of 20 homozygous mutant individuals (23X combined sequencing depth) and custom filters. Because of the lower sequencing coverage of one of the input sets we chose a lower *k, k* = 40. This approach identifies 3297 putative mutations, including 1519 calls supported by *k k*-mers each. Among these, 88% are G/C to A/T transitions, and include the ms5 causative mutation and mutations representing 10 of the key markers used in the *ms5* mapping (compare to corresponding LNISKS results in Table 2).

To provide an estimate of the specificity of the calls we have aligned pairs of postulated wild-type and mutant sequences to IWGSC RefSeq v1.0 assembly. In 2213 or 67% of cases, both sequences aligned to the same locus. Among the 1519 high confidence calls supported by *k k*-mers, 1358 (89.4%) sequence pairs aligned to the same locus. Of these, 1286 (94.7%) aligned to chromosome 3A.

Although this approach appears to be less sensitive than LNISKS, this may be at least partly due to the lower sequencing coverage (≈ 15X) available for the fertile heterozygous bulk, further exasperated by the fact that for the heterozygous portions of the genome (the region of interest), we would require twice the sequencing coverage for a comparable level of evidence to be available for calling mutations. The available ≈ 15X coverage in the region of interest translates to ≈ 7.5X available per allele.

### Availability and dependencies

The LNISKS piepline is freely available at https://github.com/rsuchecki/LNISKS under Apache 2.0 license. Many of the individual components of the pipeline are modules from https://github.com/rsuchecki/yakat toolkit (Java), which is packaged as a single executable with no dependencies. LNISKS also utilizes bash, AWK/MAWK and standard Linux tool set. It was developed and tested on Ubuntu Linux 14.04 with Java 8 update 74. It has few dependencies, with `KMC2/3` (*Deorowicz et al*., 2015; Kokot *et al*., 2017) used for *k*-mer counting and set operations and `VSEARCH` (*Rognes et al*., 2016) used for seed clustering.

## Discussion

Bulk Segregant Analysis supported by modern sequencing technology is a powerful approach for identification of mutagenesis-derived causative mutations. Our computational approach allows for it to be applied when a suitable reference genome assembly is not available. Another advantage of *k*-mer based mutation identification directly from the input data is that it is not reliant on the fine-tuning of the alignment parameters. This effectively adds to computational efficiency of the approach as it reduces the need for parameter space exploration. Where such exploration of parameter space is necessary, it is fast thanks to operating on much reduced sets of fixed length sequences (*k*-mers). Finally, as illustrated in this work, the subtraction of *k*-mers present in third party datasets amplifies the signal of the targeted mutations, thus further accelerating the process of candidate gene identification.

Low recombination around the *TaMs5-A* locus resulted in a very high number of linked mutations thus highlighting one of the main risks associated with the approach. Population size, number of meiosis events and sampling affect how many recombination events are captured for downstream analysis. This coupled with large genome size translates into considerable cost of undertaking BSA analysis. In some cases this could potentially be alleviated e.g. by selecting the most informative recombinants based on prior knowledge from genetic mapping of the locus of interest.

The IWGSC reference assembly allowed us to illustrate the validity of our approach. It also indicates that in cases where the studied accession is sufficiently similar to Chinese Spring, a reference-based approach may suffice. If however the studied cultivar has, for example, the relevant chromosome (or part of it) introgressed from a related species then an approach reliant on a reference assembly may not be suitable. Furthermore, the effective utility of this particular assembly is largely dependent on it being presented as a set of well constructed pseudo-chromosomes. Such level of contiguity and refinement was not available for any of the previous assemblies of bread wheat, and may not be available for many other species in the short to medium term.

LNISKS is developed and tested with WGS data in mind but it can be applied to other types of sequence data such as RNA-Seq. Using RNA-Seq in place of DNA-Seq offers a reduction in cost but carries certain risks, not least the risk of the targeted gene not being expressed or sufficiently highly expressed in the collected tissues at a given developmental stage. A causative mutation outside the expressed portion of the genome will not be detected directly and the identification of the relevant gene or genes by relying on changes in expression or even lack of expression carries a high level of uncertainty. On the other hand, mutations in linked genes will likely be detected and these may be used to narrow down the candidate list.

Our approach detects short indels as well as SNPs. We do not focus on this functionality, as indels are rare in EMS-derived data. For example, among category A calls at k = 54, 1 bp insertions and deletions represent 0.5% of calls, or 0.15% of calls when custom filters are applied. More generally, as already demonstrated by NIKS, also large indels can be captured, but this requires fine-tuning of parameters when pairing seeds and calling such mutations.

Finally, we have demonstrated how to identify mutations from data derived from heterozygous individuals but also how such data can be leveraged for prioritising called mutations. More generally, combining KMC for counting *k*-mers and set operations with the tool set we have developed, allows us to go beyond the paradigm of LNISKS and NIKS to identify and prioritize SNPs across multiple datasets. One could, for example discard any *k*-mers present in all the bulks/datasets under consideration and use our tool set to identify SNPs between any of the bulks.

### Experimental procedures

We provide a general overview of our approach in Figure 1. LNISKS broadly follows the steps of the established NIKS approach (Nördstrom *et al*., 2013). One of the main differences is the application of custom *k*-mer filters. Other innovations which we outline in this section pertain to extension of *k*-mers to seeds both before and after the seeds are clustered/paired. SNP prioritisation and the functional annotation of the underlying sequences are not core parts of our pipeline, but we do provide the tools required for these operations. After a brief summary of our library preparation and sequencing protocol, we outline the key elements of our pipeline, explore the issue of call prioritization and finally detail our proof-of-concept approach for reference free identification of heterozygous mutations.

### Library construction and sequencing

The genomic DNA was prepared according to a library construction protocol developed by Illumina and sequenced using the Illumina HiSeq2500. DNA was extracted from frozen tissue from 80 individual plants using the DNAeasy system (Qiagen) according to manufacturer’s conditions. Briefly, after genomic DNA was sheared by sonication using a Covaris S220/E220 system, the resulting DNA fragments were end-repaired and their 3’ ends treated for A-base addition. After ligation of Illumina-specific adapters and gel-based size-selection, adapter-ligated DNA fragments were subjected to limited PCR amplification with Illumina-specific PCR primers. Cluster generation and paired-end sequencing of the amplified DNA fragments were performed on an Illumina cBot and Illumina HiSeq2500, respectively, according to Illuminas instructions. Sequencing primer hybridization was performed on the cBot and 151 cycle paired-end protocols were used on the HiSeq2500. Sequences and quality scores were generated with the Illumina pipeline software for image analysis and base calling. After initial base calling and processing, the sequencing files generated by the Illumina pipeline were converted to FASTQ format and additional custom quality filtering was performed, such that reads were trimmed if they harboured one or more base at their 3’ end with a quality score ¡ 15. Assuming 17 Gbp genome size, sequencing coverage for the bulks was ≈ 19X for the 20 homozygous wild-type plants, ≈ 23X for the 40 homozygous mutant plants and ≈ 15X for the additional 20 wild-type plants heterozygous for the locus.

### Custom k-mer filters

We have devised a filtering strategy which reduces the computational cost of extending and clustering seeds, and reduces the number of false positive calls. The mode of use as well as usefulness of such filters depends on *a priori* knowledge and the availability of suitable sequence data. Filters are in principle best suited for aiding the detection of dominant mutations when sufficient sequence and phenotypic data are available. When looking for a dominant allele responsible for a trait we do not expect its exact sequence to be present in a variety not displaying that trait. In the case of *ms5* the assumption is that a mutation causing such an unambiguous phenotype (as male sterility) should not be present in cultivated varieties, so the application of a filter was straightforward also for this recessive mutation. Consequently, we are able to extract sets of filtering *k*-mers from datasets such as the WGS data of 16, predominantly Australian wheat cultivars (Edwards *et al*., 2012). We used KMC2 to extract *k*-mers, k G {24, 32,40,48, 52, 54, 56, 58, 60, 64, 72} from each of the 16 datasets. For each k we have computed a union of the 16 sets, only considering *k*-mers occurring at least twice in a given set, *k*-mers occurring only once in a dataset are ignored as they are likely to arise from sequencing errors in the input reads. Each set F generated this way holds from 10. 2 × 10^9^ to 16.8 × 10^9^ *k*-mers with the corresponding KMC database occupying from 41 gigabytes for k = 24 to 338 gigabytes for *k* = 72, or the total of 2.2 terabytes for the explored values of k. We expect that LNISKS user would only need to generate a database for a single selected k value. The relations between a filtering set and the input sets of *k*-mers for the selected value *k* = 54 are illustrated in Figure 5.

**Figure 5:**
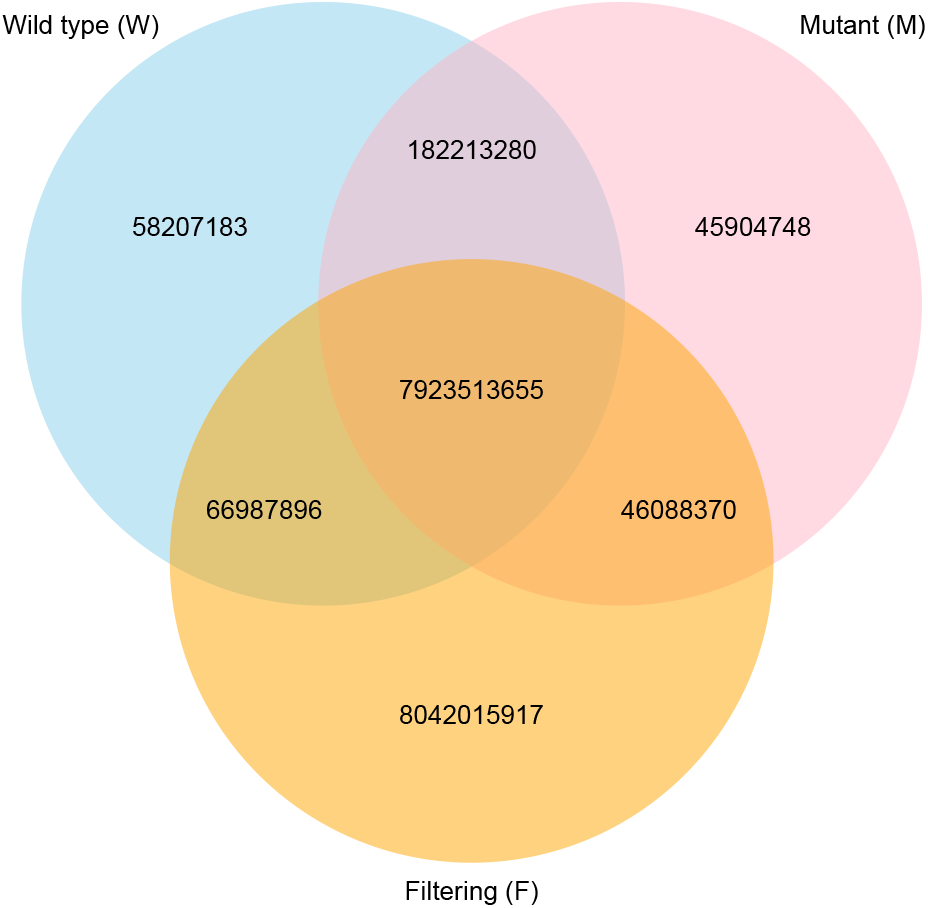
Relations between sets of 54-mers occurring at least twice in each of the datasets. The discarding of *W* ∩ *M* greatly reduces computational complexity. This operation discards 8105 726 935 *k*-mers and results in the set *M′* = *M* \ *W*, |*M′*| = 91 993 118. In addition, the application of custom filtering set *F* allows us to half the number of the *k*-mers remaining in *M′* as we discard *F* ∩ *M′* i.e. 46 088 370 *k*-mers. The value of the filtering comes from the resulting reduction in the number of non-EMS mutations called.

### Generation of unambiguous *k*-mer extensions

We have developed a specialized tool for efficient generation of unambiguous extensions (unitigs) from sets of *k*-mers. Our kextender exploits the fact that due to only sample-specific *k*-mers being present in the input, the implicit De Bruijn graph consists largely, although initially not exclusively, of disconnected sub-graphs, each representing a unitig. As we construct a map which stores the information from the input *k*-mers, we ensure that this holds for all sub-graphs. For each input *k*-mer we store two (*k* – 1)-mers, each with a single bp extension representing the first and the *k*th base, respectively. Binary encoded canonical representation of a (*k* – 1)-mer forms the key for a given entry. With each key, we can associate up to four alternative 1 bp extensions on each of the two sides. Presence of *k*-mers which are adjacent in the original input sequence limits the overhead of storing two (*k* – 1)-mers rather than a single *k*-mer to typically less than 10%. At this modest cost we gain a convenient, implicit representation of the De Bruin graph, well suited for parallel traversal for contig construction. A collision in the map construction occurs when for a given (*k* – 1)-mer we store an alternative 1 bp extension at either of the two positions – this invalidates the map entry for that (*k* – 1)-mer. After the construction stage we purge the invalidated entries from the map, which ensures that the underlying implicit De Bruijn graph only contains nodes of degree ≤ 2. Any entry which only holds a single 1 bp extension is also purged as it only holds redundant information which is also stored with another (*k* – 1)-mer. At this stage, map entries representing graph nodes adjacent to the purged ones are recorded in set *T*. Sub-sets of *T* are then passed to extender threads which traverse the sub-graphs associated with each element independently in parallel to build contigs. If two threads happen to start traversing a path underlying the same unitig from opposite ends, one thread abandons the extension, leaving the other one to continue the extension process unhindered. Such collisions are rare, at 1 per 25 000 extensions in our experimental set-up. This will vary depending on size and contiguity of the graph being traversed as well as the number of threads traversing it.

### Further extension of clustered seeds

In the NIKS approach, seeds from the two datasets are paired based on sequence similarity. In principle this should suffice. In practice however, *k*-mer extension to seeds may often fail to reach the desired length of 2*k* – 1 bp. This can be due to insufficient or uneven sequencing coverage or repetitiveness of the sequence in question, which translates to non-uniqueness of some of the *k*-mers which overlap a putative mutation. The issue of non-uniqueness of *k*-mers can be alleviated by increasing *k*. However as we increase *k*, seed contiguity may suffer due to insufficient or uneven coverage. Because of that, rather than simply finding best matching pairs of wild-type and mutant sequences we have opted for similarity-based clustering of all sequences from the two input datasets. If both the wild-type and the mutant sequence are 2*k* – 1 bp they will most likely be clustered together as they would if we were simply pairing sequences across the two sets. Clustering however also allows us to group together multiple incompletely extended seeds which overlap a putative mutation site. Our vclusters module prior to variant calling merges clustered contigs from a given set. For that we exploit the fact that wild-type and mutant contigs clustered together serve as reciprocal anchors thereby facilitating extension of contigs to 2*k* – 1 bp. This enables us to identify mutations which would otherwise remain undetected or classified at a low confidence level among mostly false calls. This extension procedure is only performed if there are no mismatches between the contigs within a given data set. This subroutine is behind ≈ 20% of category A calls across the *k*-mer lengths explored in our experiments.

### Call prioritisation through *k*-mer threading

We developed the snpmers module which facilitates rapid, reference- and alignment-free genotyping of SGS datasets for a pre-defined set of varying loci, such as the set of category A calls from LNISKS. Information generated by this module also allows us to prioritize candidate mutation calls. In addition to the homozygous wild-type (*Ms5*) and homozygous mutant (*ms5*) data, we can also tap into the respective heterozygous wild-type data. We use the last of these datasets to illustrate the workings of snpmers but the module’s application is in no way limited to heterozygous data.

We start by extracting set H of *k*-mers from the WGS reads obtained from fertile plants heterozygous for the *ms5* mutation at the *TaMs5-A* locus. To speed-up the subsequent steps, we start by discarding *k*-mers which are present in both the wild-type and the mutant as these do not capture any of the mutations. We use KMC2/3 to subtract the intersection of those two sets from *H*, i.e. *H′* ← *H* \ (*W* ∩ *M*). We next employ our genotyping module snpmers which takes the list of variants called so far as well as a set of *k*-mers (in this case, *H′*). Let 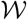 be the set of all possible *k*-mers overlapping a variant site and matching the wild-type allele, and let 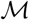 be the set of all possible *k*-mers overlapping a variant site and matching the mutant allele. The *k*-mers from *H′* are matched to a variant position and their frequencies are recorded, resulting in sets 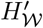 and 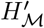 of *k*-mers from *H′* which match the wild-type and the mutant allele, respectively. This provides us with two measures of support for a given allele at a position:

i. the median frequency of *k*-mers in 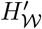 (or 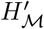)
ii. the *k*-mer coverage ratio of a given allele 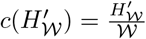 (or similarly 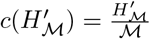)

Based on these measures, the snpmers module genotypes the input sample for each of the input loci. In the context of *TaMs5-A* this allows us to prioritize original calls which are also called as heterozygous in *H*.

More generally, this approach allows us to assign the two aforementioned measures to both alleles of each locus for each of our input datasets, including *W* and *M*. These can then be used to sort or filter the list of putative mutations to be able to focus on the highest confidence calls. The same measures can also be computed for individuals which constitute the bulks. Due to shallow sequencing of individual plants we cannot draw conclusions from absence of evidence for a given allele in a single plant, but wherever there are *k*-mers supporting or contradicting a given call, these can be quantified to approximate the number of plants which support or contradict that call. For a given call we record how many of the mutant plants yielded one or more *k*-mers supporting the postulated mutant allele. We call these mutant-true (MTT). We also record how many of the mutant plants yielded one or more *k*-mers supporting the postulated wild-type allele, i.e. mutant-false (MTF). Similarly, we establish the wild-type true (WTT) and wild-type false (WTF) values. Finally we define the support value for a given call by adding the supporting values and subtracting the contradicting values, i.e. *Support = MTT + WTT – MTF – WTF*. Weighing can be applied to highlight the the presence of *k*-mers contradicting the expected allele, as apart from error or repeat-derived contradicting *k*-mers, these are an indication of a given locus not being strongly linked to the causative one and consequently different in some of the individuals in the same bulk. Alternatively, MTT and WTT values can be considered separately form MTF and WTF. The sum of MTF and WTF is the the evidence contradicting a particular mutation call, and as illustrated by Figure 6, it is least pronounced in the centromeric block which contains the *ms5* mutation.

**Figure 6:**
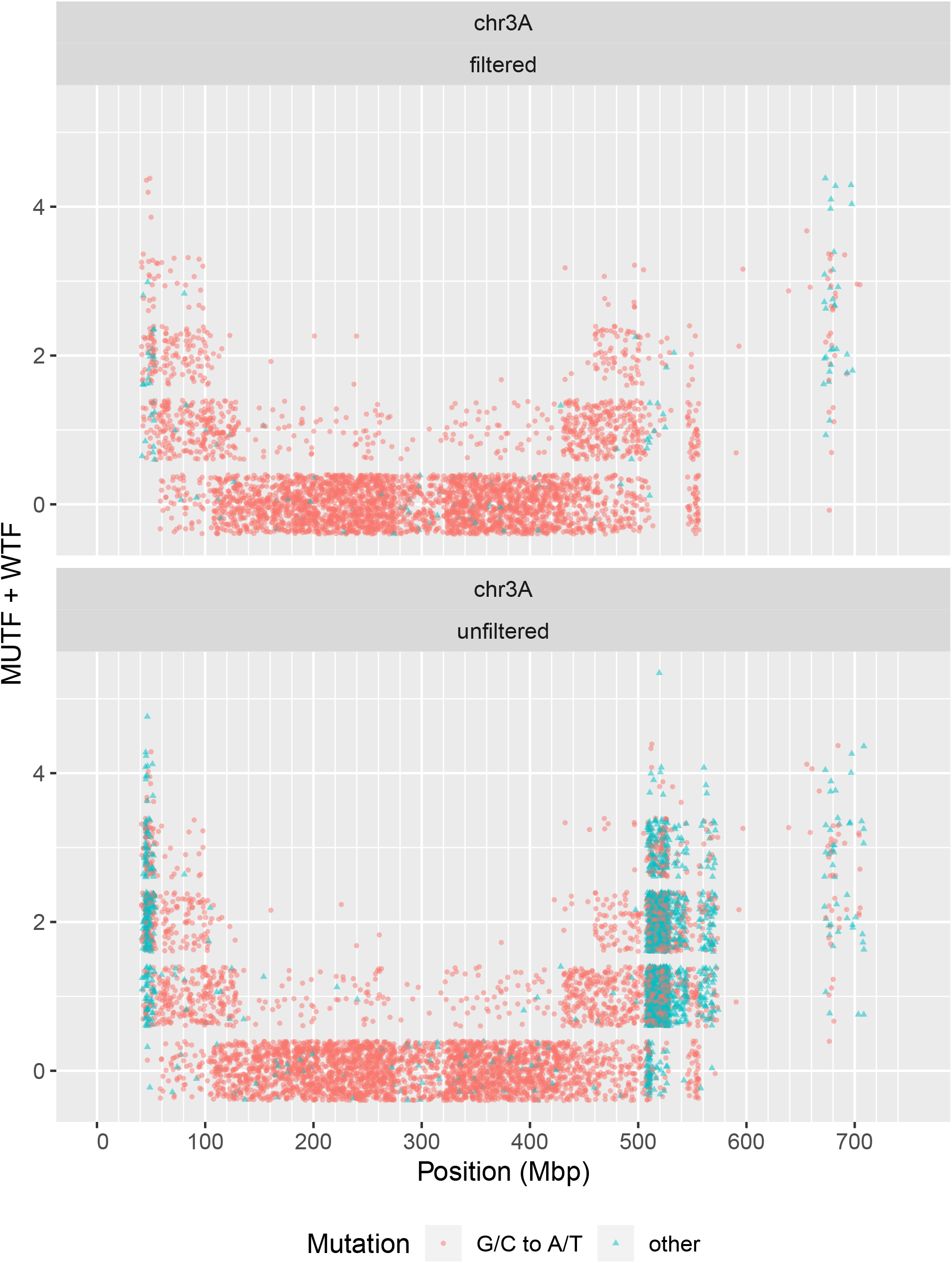
Identified category A mutations distributed along the 3A pseudo-chromosome. Towards the telomeres, an increasing proportion of calls are contradicted by *k*-mers from an increasing number of individuals.

### Proof-of-concept procedure for identification of heterozygous mutations

LNISKS (and NIKS) are designed for identification of homozygous mutations. Here we outline a procedure for the identification of heterozygous mutations using the LNISKS tool set. We first subtract the set *M* of *ms5* mutant *k*-mers from the set *H* of *Ms5* heterozygous fertile *k*-mers and subtract the set *F* of custom filtering *k*-mers from the *ms5* mutant data set *M*, that is: *H* ← *H* \ *M* and *Mf* ← *M* \ *F*. We then extend the *k*-mers from *H′* to obtain the set of unitigs (or seeds) representing the wild-type genotype. We only keep seeds of length 2*k* – 1 and assume that these harbour mutations at the kth base. We employ our seedmers module to match the mutant *k*-mers from *M_f_* to the seeds, allowing a mismatch at the *k*th base. For each seed we record frequencies of up to *k k*-mers representing an allele alternative to the wild-type represented by the seed. The collected information forms the bases of genotype calls.

### Long unpaired seeds

Sequences above user defined length which remain unpaired are output by our pipeline and in some scenarios may be of interest as these may reflect e.g. presence of larger indels or sequences highly divergent between the studied datasets. In other cases this output is more likely to represent an assembly of contaminant sequences.

## Acknowledgements

This research was funded by DuPont Pioneer Hi-Bred International Inc. We are grateful for additional support provided by The University of Adelaide. The authors thank the International Wheat Genome Sequencing Consortium (http://www.wheatgenome.org) for providing pre-publication access to IWGSC RefSeq v1.0.

